# Chemogenetic inhibition of the noradrenergic locus coeruleus promotes the development of risk-taking decisional strategies and selectively enhances motor impulsivity in females

**DOI:** 10.64898/2026.07.02.736138

**Authors:** Chloe S. Chernoff, Tristan J. Hynes, Dimitrios K. Avramidis, Shrishti Ramaiah, Amanda C. Lee, Aryan Khoshnevis, Kelly M. Hrelja, Catharine A. Winstanley

**Affiliations:** Graduate Program in Neuroscience, Department of Psychology, Djavad Mowafaghian Centre for Brain Health, University of British Columbia, Vancouver, Canada, V6T 1Z3; Department of Psychology, Djavad Mowafaghian Centre for Brain Health, University of British Columbia, Vancouver, BC, Canada, V6T 1Z4

**Keywords:** noradrenaline, locus coeruleus, impulsivity, decision making, sex differences

## Abstract

The locus coeruleus noradrenaline (LC-NA) system is a key regulator of arousal, attention, and reward learning. Noradrenaline plays a critical role in impulse control, and recent evidence indicates the importance of noradrenaline signaling in cost-benefit decision making once choice strategies are established. However, whether the LC causally shapes the acquisition of decision strategies, and how this contribution may differ across sexes, remains unclear. We addressed these questions by chemogenetically inhibiting catecholaminergic neurons within the LC of adult tyrosine-hydroxylase Cre (TH::Cre) rats (n=69; 35 females) throughout acquisition of the cued rat gambling task (crGT), a probabilistic decision making paradigm that incorporates salient audiovisual reward-paired cues and simultaneously measures motor impulsivity. LC inhibition accelerated the development of risky choice strategies early in training in both males and females, reflected by impaired adoption of the most advantageous option and increased preference for risky options. Trial-by-trial analyses reveal that LC inhibition promoted switches in choice strategy following safe wins, while reducing switches away from risky options after both wins and losses. LC inhibition therefore seemed to encourage the repetition of actions that resulted in more uncertain outcomes. LC inhibition also selectively enhanced motor impulsivity in females, particularly early in training. These results provide causal evidence that the LC system guides the formation of optimal decisional strategies, while exerting sex-specific control over impulsive action.

**Abbreviated summary:** Chernoff et al. demonstrate that noradrenergic locus coeruleus (LC) neurons are required to develop optimal decisional strategies, as their inhibition hastens the adoption of risk-preferring choice tendencies. This may be underpinned by reduced behavioural sensitivity to likely outcomes. They also show that the LC regulates impulse control, specifically in females.

## Introduction

The locus coeruleus noradrenaline (LC-NA) system modulates arousal and cognition to regulate behaviour (for review: Sara & Bouret, 2012). Multiple lines of evidence indicate that the LC mediates switches between ‘exploitation’ of a lucrative strategy and ‘exploration’ of other potentially fruitful options when the utility of the active strategy wanes (Aston-Jones & Cohen, 2005; Bouret & Sara, 2004; Jordan, 2024; Sales et al., 2019; Tervo et al., 2014). The LC may encode reward prediction errors (RPEs) and drive adaptive behaviour through RPE-based reinforcement learning (Breton-Provencher et al., 2022; Carvalheiro & Philiastides, 2023; Su & Cohen, 2022). The LC also responds to salient or motivationally-relevant stimuli (Aston-Jones et al., 1994; Aston-Jones & Bloom, 1981; Bouret & Richmond, 2015; Grant et al., 1988; Rajkowski et al., 2004; Vazey et al., 2018) and is required for learning stimulus-guided behaviours (Selden, Cole, et al., 1990; Selden, Robbins, et al., 1990), altogether situating the LC as an orchestrator of cue-guided behaviour.

Recent evidence highlights that noradrenaline’s impact on risk-taking depends on the presence of salient reward cues. In the cued rat gambling task (crGT), rats choose among four options with differing probabilities of sugar pellet rewards and timeout punishments, where the optimal strategy is to favor the low-risk/low-reward options that maximize overall gains (Barrus & Winstanley, 2016). Noradrenergic compounds consistently bias choice toward these safer, more lucrative options, but only when audiovisual cues accompany rewards (Baarendse et al., 2014; Chernoff et al., 2021, 2024; Silveira et al., 2016). This cue-specific effect aligns with the LC’s sensitivity to salient stimuli. Importantly, these findings show that elevating noradrenaline can reshape already-established strategies, leaving open the question of how LC–NA activity shapes the *acquisition* of decision strategies on the crGT.

Manipulations of noradrenaline potently regulate motor impulsivity across various preclinical assays, including the rGT) (Baarendse et al., 2014; Chernoff et al., 2021, 2024; Silveira et al., 2016), which measures motor impulsivity simultaneous to decision making in a manner akin to the 5-choice serial reaction time task (5CSRTT (Benn & Robinson, 2017; Economidou et al., 2012; Navarra et al., 2008; Robinson et al., 2008). Functional alterations in the noradrenaline system may also underpin interindividual differences in trait impulsivity and vulnerability to impulse control disorders (Ansquer et al., 2014; Chernoff et al., 2025; Engberg et al., 1987; Hesse et al., 2017; Russell et al., 2000; Whelan et al., 2012; Williams & Potenza, 2008). Further, the LC-NA system exhibits prominent morphological (Bangasser et al., 2011; Pinos et al., 2001), molecular (Bangasser et al., 2010, 2013), transcriptional (Mulvey et al., 2018), and electrophysiological (Bangasser et al., 2013; Curtis et al., 2006; Mariscal et al., 2023) sex-differences which may render females more responsive to stressors or high arousal states (Bangasser et al., 2019). Despite strong evidence for distinct LC-NA system characteristics in males and females, it remains unclear how these differences may manifest to guide probabilistic decision making in a sex-dependent manner.

Given that LC-NA facilitates adaptive action selection by leveraging environmental cues through attentional and RPE-based processes, we hypothesized that the LC plays a causal role in the development of decisional tendencies throughout acquisition of the crGT. We also predicted that LC-NA may more markedly influence behaviour in females, due to the greater sensitivity of the female LC-NA system. We tested these hypotheses by employing chemogenetics to selectively inhibit catecholaminergic – i.e., putative noradrenergic – cells of the LC during the acquisition of the crGT in adult male and female rats. We used trial-by-trial analyses to explore whether any observed changes in decision making following LC inhibition could be driven by alterations in behavioural sensitivity to wins or losses.

## Methods

### Subjects

Sixty-nine adult TH::Cre rats (34 males, 35 females) were used for the behavioural experiments described below, with the experiment run in two roughly equally sized independent cohorts (n=32; n=37). Rats were bred in house on a Long Evans background strain (Charles Rivers Laboratories, St. Constant, QC, Canada) expressing a Cre recombinase enzyme in tyrosine hydroxylase (TH) expressing neurons (TH::Cre) (Long-Evans-TG (TH-Cre)3.1Deis, RRRC #00659; Rat Resource and Research Centre, RRRC, Columbia, MO, USA), and were 10-12 weeks of age at the beginning of the experiment. Rats were randomly assigned to the experimental group, which would receive bilateral infusion of a Cre-dependent virus to express the inhibitory designer receptor (DREADD), hM4 (AAV2/9-hSyn-DIO-hM4(Di)-mCherry; #44362-AAV9, AddGene, MA, USA), in the LC, or to the control group. The control group was comprised of an equal number of transgene positive (TG+) and transgene negative (TG-) rats, which all received intra-LC infusions of a Cre-dependent virus carrying the construct for the control fluorophore, mCherry (AAV2/9-hSyn-DIO-mCherry; #50459-AAV9, AddGene, MA, USA). Post-surgery group totals were as follows: 18 hM4 males, 16 control males, 17 hM4 females, 18 control females. All animals were housed with littermates in same-sex pairs or trios and kept in a climate-controlled colony room on a reverse 12-h light-dark cycle (lights off at 08:00; temperature at 21°C ±2°C). Water was offered *ad libitum* in the home cage for the entirety of the study and the animals were maintained on standard rat chow in addition to the sugar pellets (∼5g/day) earned in the task. All housing conditions and testing procedures were in accordance with the guidelines of the Canadian Council on Animal Care and all protocols were approved by the Animal Care Committee of the University of British Columbia.

### Experimental timeline

A schematic summary of the experimental timeline is provided in Figure 1A. First, rats underwent stereotaxic surgery followed by a period of at least four weeks to allow for post-operative recovery and ample viral-mediated expression of the inhibitory DREADD or control fluorophore in TH+ LC neurons before behavioural training began. At least one week before beginning training on the crGT, the rats were food restricted to ∼85% of their free feeding weight (∼15 g of standard rat chow/day for males and ∼11g/day for females). Throughout the study, each rat underwent behavioural training in the same operant chamber, and at the same time of day (11:30-12:00 or 12:00-12:30).

**Figure 1:**
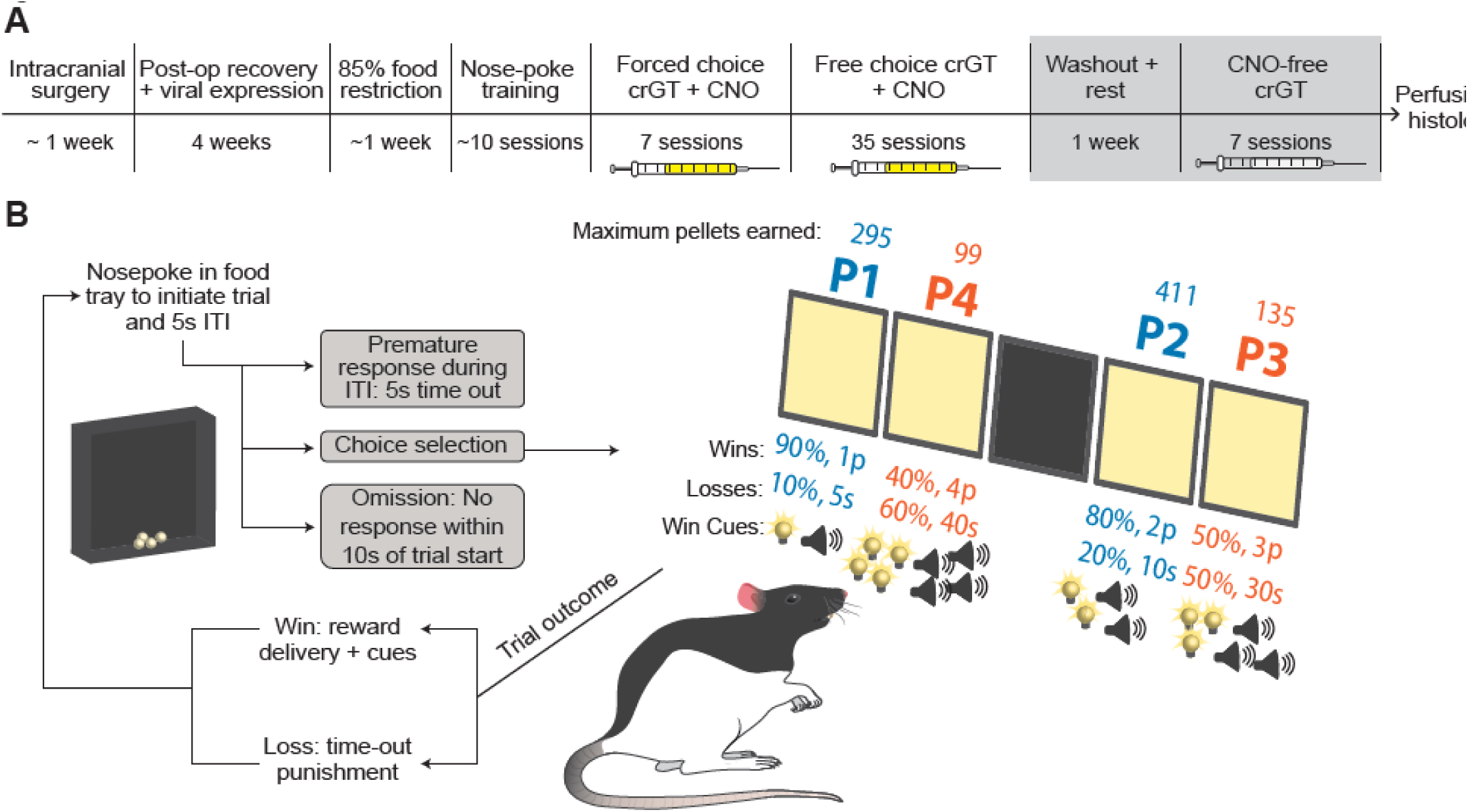
Experimental timeline and cued rat gambling task (crGT) schematic. **A)** The experiment began with intracranial surgery to infuse a Cre-dependent viral vector into the LC of male and female tyrosine hydroxylase (TH)::Cre rats to express either an inhibitory designer receptor (DREADD), hM4(Di), or a control fluorophore, mCherry. Controls were an equal number of transgene positive (TG+) and negative (TG-) animals. Following post-operative recovery and at least 3 weeks to allow for viral expression, rats were food restricted to 85-90% body weight, habituated to reward pellets and the operant chambers, then trained to make nose-poke responses at the aperture of the five hole array for food. Next rats were exposed to task contingencies through 7 days of forced choice training, before moving on to 35 days of free choice crGT whereby all four options were available to chose from. Syringe icons indicate pre-session injections of CNO (yellow; 1mg/kg i.p.) or saline (grey; 1mL/kg i.p.) 30 mins before the task. Following the CNO-concurrent free choice crGT, rats were given one week for CNO to washout, and were re-tested CNO-free for 7 sessions of free choice crGT. After behavioural testing was complete, rats were transcardially perfused, and brains were extracted and processed for immunohistochemical confirmation of DREADD expression. **B)** A schematic representation of crGT task structure and probabilistic reward contingencies of all four options. A trial begins with an initiating nosepoke into the food magazine opposite the 5-hole array. After a five second inter-trial interval (ITI), all four options are illuminated on the 5-hole array (middle hole is not used in the crGT) and a rat can nosepoke into any hole to make a choice. Each option is associated with a distinct probability and magnitude of sugar pellet wins and time out punishment such that safe options (P1&P2) yield smaller more likely wins and shorter less likely punishments than risky option (P3&P4). Win-paired audiovisual cues accompany reward delivery on winning trials, and scale in complexity with the magnitude of wins (ie. riskiest options are associated with the most complex cues). Motor ismpulsivity is assessed by the rate at which rats prematurely respond at an option prior to the completion of the 5s ITI.

Behavioural training was conducted as described previously (Barrus & Winstanley, 2016, see below for further details). Thirty minutes prior to each session during both the forced choice and free choice segments of crGT training, rats received intraperitoneal (IP) injections of clozapine-N-oxide (CNO) in 5% DMSO (1 mg/kg at a volume of 1mL/kg). After the completion of 35 free choice crGT sessions with pre-session CNO injections, rats underwent one week of CNO washout, during which no behavioural testing occurred, then completed an additional seven CNO-free crGT sessions identical to the free choice crGT acquisition sessions except that pre-session injections were 0.9% saline (1ml/kg i.p.) instead of CNO. Following all behavioural testing, rats were euthanized via transcardial perfusion with 0.01M PBS and 4% neutral buffered formalin (NBF), and the brains were extracted and processed for immunohistochemistry.

### Stereotaxic surgery

Subjects were induced under 5% isoflurane in oxygen at 2L/min before being placed into a stereotaxic frame with the skull flat (bregma and lambda at the same DV coordinate). A surgical plane of anesthesia was maintained at 2% isoflurane in oxygen. The stereotax arm was angled 15 degrees backward in the AP plane to avoid the transverse sinus. Each rat received bilateral 1.2 μL infusions of a Cre-dependent virus into the LC (at a 15° angle- AP: −11.6 from bregma, ML: +/− 1.3, DV: −6.9 from skull surface), via two 31G injectors and at a rate of 0.2 μL/min, to express either an inhibitory designer receptor (DREADD) (AAV2/9-hSyn-DIO-hM4(Di)-mCherry; Addgene plasmid # 44362) in experimental rats or control fluorophore (AAV2/9-hSyn-DIO-mCherry; Addgene plasmid # 50459) in TG+ controls (Figure 2A). TG-controls received the same Cre-dependent mCherry virus as TG+ controls, yet would not express the control fluorophore given the absence of Cre-recombinase. After the infusion was complete, the injector tip was left in place for a 5 minute diffusion period.

**Figure 2:**
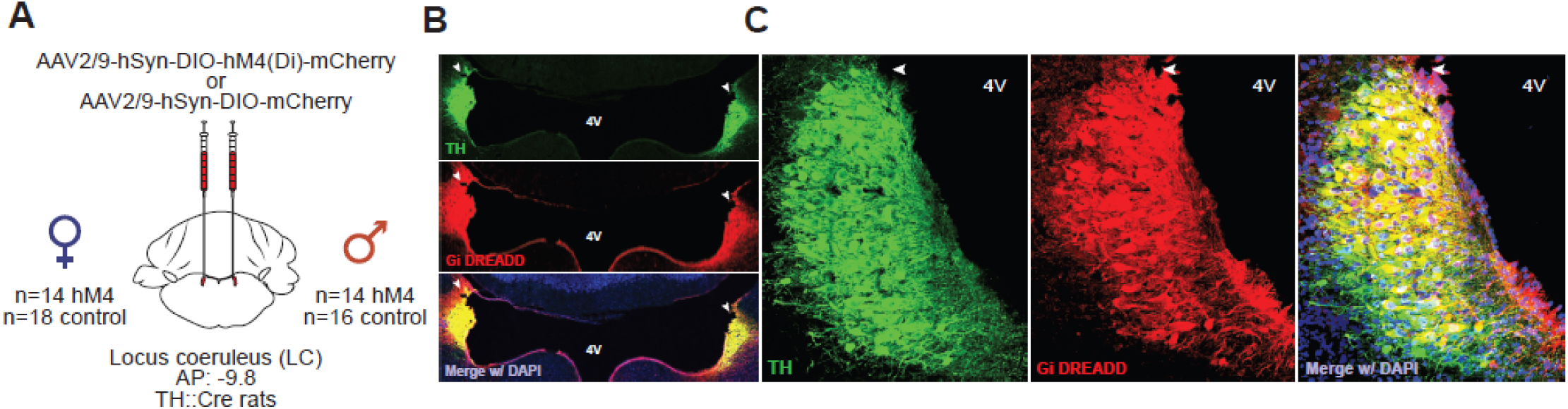
Immunohistochemical confirmation of expression of an inhibitory DREADD in TH-positive cells of the locus coeruleus (LC). **A)** An illustration of the viral preparation for inducing DREADD or control fluorophore expression in TH+ LC cells, including viral vectors and final sample size following histological exclusion due to inadequate DREADD expression. **B)** Confocal micrographs at 20x magnification depicting a representative image of bilateral hM4 DREADD expression (red) in TH+ (green) cells of the LC. **C)** A 63x magnification confocal image of colocalized TH (green) and hM4 DREADD (red) expression in the LC. Green = TH. Red = mCherry tag on hM4 DREADD. Blue = DAPI (nuclear marker). Yellow indicates co-expression of TH and the DREADD. 4V = fourth ventricle. Infusion sites are indicated with a white arrowhead.

### Apparatus

Behavioural testing took place in two adjacent rooms each containing 16 identical operant chambers (30.5 × 24 × 21 cm; Med Associates, St. Albans, VT, USA). Each chamber was enclosed in a separate ventilated sound-attenuating cabinet (Med Associates, St. Albans, VT, USA), and fans were added for increased ventilation and noise cancellation. The boxes had a curved wall on one end, fitted with an array of five nose-poke holes. Each hole contained an infrared detector and a yellow LED stimulus light. Dustless sucrose-covered food pellets (45 mg, Formula P; Bio-Serv, Frenchtown, NJ, USA) were delivered from a dispenser into a food hopper located on the opposite wall. To illuminate the chamber, a white hooded house light was located on the chamber ceiling. Control of the apparatus and data collection was performed using code written by CAW in MEDPC (Med Associates) running on standard IBM-compatible computers.

### The cued rat gambling task (crGT)

All rats underwent basic nosepoke and crGT training as described previously (Barrus & Winstanley, 2016) and illustrated in Figure 1B. Briefly, the rats were first habituated to the operant chambers in single 30-minute exposures on 2 consecutive days. All lights were turned off, and sugar pellets were placed in each of the chamber’s response holes to motivate the animals to explore fully. Habituation was considered complete once the animals reliably consumed all pellets. The animals were then encouraged to nose-poke only in illuminated holes along the curved wall using a protocol adapted from the 5-choice serial reaction time task (5CSRTT) training. Once subjects passed criterion for nose-poke training, they moved on to forced choice crGT sessions. In the forced choice crGT, each nose-poke hole was associated with varying sugar pellet rewards, audiovisual cues, and time-out punishments analogous to those of the real crGT. The rats were presented with one illuminated option at a time, with roughly equal numbers of discrete presentations of each option, such that the animals had equivalent experience with all task contingencies. The rats underwent seven sessions of forced choice training before moving on to the free choice crGT.

The procedure for the crGT was identical to that of the forced choice crGT, except all four options (P1-P4) were illuminated and available for rats to choose on each trial. Between rats, options were counterbalanced across holes to account for potential side biases. Briefly, the crGT began after a subject initiated a trial with a nose-poke in the illuminated food tray, which then began the 5-second intertrial interval (ITI). A nose-poke in any hole during the ITI was recorded as a premature response and punished with a 5-second time out with the house light on, during which no responses can be made. If the rats did not respond prematurely, stimulus lights were turned on for all four active nosepoke holes in the array, and a choice could be made. A nose-poke in one of the active holes resulted in all stimulus lights turning off and the delivery of a reward or punishment according to the reinforcement schedule of the chosen hole. Sugar pellet rewards on winning trials were delivered to the food hopper and accompanied by various patterns of audio tones and flashing stimulus lights. The audiovisual win cues scaled in intensity and complexity with the size of the reward, and therefore the ‘riskiness’ of the option (Barrus & Winstanley, 2016). On losing trials, animals were punished by a time-out of varying duration. Riskier options were paired with longer and more frequent time-out punishments. An omission was recorded if, after initiation of a trial, the rat failed to respond at any active nosepoke hole within 10 seconds of the stimulus lights coming on.

### Histology

In order to visualize hM4 DREADD expression in LC neurons, we double-labelled 35-μm sections coronal sections with primary antibodies against mCherry (Cat#ab205402; Abcam; Toronto, ON, Canada; 1:500) and tyrosine hydroxylase (TH) (Cat#AB152; Millipore Sigma; Oakville, ON, Canada; 1:500) for 32 h. Sections were then washed in PBS and incubated with secondary antibodies conjugated to Alexa Fluor® 488 (Cat#A-21103) and Alexa Fluor® 633 (Cat#A-11034) (Thermo Fischer Scientific; Burnaby, BC, Canada; 1:500 for both), followed by DAPI (Cat# 508741, 1:1000, Millipore, Etobicoke, ON, Canada). Sections were then cover-slipped under HARLECO® Krystalon™ mounting medium (Thermo Fischer Scientific; Burnaby, BC, Canada) and visualized using a SP8 WLL confocal microscope (Leica Microsystems, Germany). Subjects were excluded from statistical analyses if they failed to sufficiently express the DREADD in LC TH neurons, as evidenced by little to no mCherry and TH co-staining in the LC, or if they expressed significant ectopic expression indicated by prominent mCherry expression in non-TH positive cells. A representative image of acceptable LC hM4 expression is provided in Figure 2B,C. Following post-histological exclusion, 14 hM4 males, 16 control males, 14 hM4 females, and 18 control females were included in subsequent statistical analyses.

### Statistical analysis

The following behavioural crGT variables were analyzed for each subject on each session: decision making score calculated based on the proportional number of choices made of each safe and risky option [(P1 + P2) − (P3 + P4)], percent choice of each option (number of PX chosen/total number of choices × 100), percentage of premature responses (number of premature responses/total number of trials initiated × 100), sum of omitted responses, sum of trials completed, and average latencies to choose an option and collect reward.

#### Analysis of variance (ANOVA)

The effect of our chemogenetic manipulation on each behavioural variable was assessed using a repeated measures ANOVA performed across CNO crGT acquisition sessions with sex (male, female) and group (hM4, control) as between-subject factors. crGT sessions were split into three blocks to assess the impact of chemogenetic LC inhibition across different epochs of task acquisition, given that the NA system may have unique influences on the early acquisition of behaviour compared to its expression once well-established (Ansquer et al., 2014; Chernoff et al., 2025). Session and block were included as within-subject factors for all repeated measured ANOVAs. Following significant session-wise interactions, group means from each session were subject to post hoc planned comparison contrasts between hM4 and controls.

#### Regression trajectory analyses across acquisition

To describe and compare trajectories of behaviour across acquisition between hM4 and control groups, behavioural variables that resulted in significant group-wise within-subject contrasts were fit with linear or non-linear regression lines, depending on the polynomial order of the significant contrast. If more than one polynomial order was significant for a given contrast in the ANOVA, the best fitting regression equation was determined based on the model that returned the lowest RMSE value and confirmed using an extra sum-of-squares F-test with the simplest (ie. lowest polynomial) model as the null hypothesis. To investigate potential differences in behavioural trajectories of hM4 and control rats, the best fitting regression equations for experimental and control groups were compared using an F-test to determine if one equation could adequately fit both subsets of data. If one equation was unable to fit both sets of data, the unshared coefficients of each function were compared with an F-test to assess which element differed between each function of best fit. The following coefficients were considered: B0 (intercept), B1 (linear constant), B2 (quadratic constant), and B3 (cubic constant) (Equation 1).

**Equation 1**. *Coefficient naming conventions for regression lines/curves of best fit.* Example equation for A) linear, B) quadratic, and C) cubic functions, where X = crGT session.

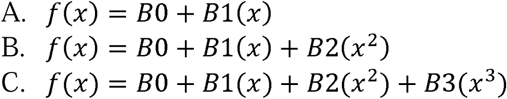

#### Win-stay/lose-shift (WSLS) analyses

Trial-by-trial data were used to assess how LC inhibition may alter the way in which a given outcome influences choice on the following trial, and whether LC manipulation may preferentially alter the way in which wins or losses inform subsequent decisions. For each rat, the outcome of every trial was identified as either a win or loss from the safe (P1 or P2) or risky (P3 or P4) choice category. The choice made on the following trial was deemed a ‘shift’ if the rat chose an option from the other category [ie. safe (P2) ➔ risky (P4)]. The proportion of trials upon which choice shifted was calculated for each type of preceding outcome: safe win, safe loss, risky win, or risky loss. Proportion data were analyzed using repeated measures ANOVA and planned comparisons as other behavioural variables, detailed above.

SPSS Statistics 28.0 (IBM, Chicago, IL, USA) was used to conduct all ANOVA and pairwise analyses. Linear and non-linear regression analyses were performed using GraphPad Prism 10.0 (GraphPad, Boston, MA, USA). Trial-by-trial variables for win-stay/lose-shift analyses were calculated using a custom Python code, and analyzed in SPSS. Effect sizes are reported as partial eta squared (ηp^2^) and Cohen’s d for ANOVA and planned comparison results, respectively.

## Results

### Inhibitory DREADD expression is selective to TH+ cells of the LC

Six hM4 rats (2 female, 4 male) were excluded due to an absence or near absence of mCherry-tagged DREADD in TH-positive LC cells. All other rats exhibited ample viral expression of the inhibitory DREADD in TH positive cells (Figure 2B,C). One hM4 female was euthanized during crGT training due to health reasons unrelated to chemogenetic inhibition of the LC. Following post-histological exclusions, 18 control females, 16 control males, 14 hM4 females and 14 hM4 males were included in the statistical analyses (Figure 2A).

### Cre-dependent expression of a control fluorophore in the LC does not influence crGT performance

The expression of an mCherry control fluorophore in the LC and Cre recombinase in TH positive neurons did not influence performance on the crGT as TG- and mCherry-expressing TG+ rats did not differ on any crGT variable across the entirety of the experiment (Supplemental Figure 1A-J; controls- effect of virus: all Fs≤1.914, ps≥0.177). Thus, TG+ and TG- rats were pooled as controls.

### LC inhibition promotes risk-taking during crGT acquisition

Chemogenetic inhibition of catecholaminergic LC neurons during acquisition of the crGT significantly impaired decision making (Figure 3A; block x session x group: F_22,1012_=2.247, p=0.021; ηp^2^=0.040). Specifically, hM4 animals had lower decision making scores than controls on sessions 9, 11 and 12 (Figure 3A; hM4 vs controls: all ps≤0.045, ds≥0.522; other sessions: ps≥0.068), indicating riskier decision making that was restricted to the early to middle stages of task acquisition. The line of best fit for decision making score also differed between groups (block x session x group- linear contrast: F_1,54_=5.297, p=0.025, ηp^2^=0.089; quadratic contrast: F_1,54_=5.377, p=0.024, ηp^2^=0.091) whereby the trajectory of score for the hM4 group was best fit with a quadratic equation (linear fit: F_1,977_=13.80, p<0.001) while a linear model best fit control data (linear fit: F_1,187_=2.045, p=0.153). The resulting equations of best fit indicate that both groups exhibited a decline in score across training, as consistently observed in the crGT (Barrus & Winstanley, 2016; Hathaway et al., 2024; Mortazavi et al., 2023), yet the hM4 group experienced a steeper decrease in score during early to middle acquisition compared to control rats. No interactions with sex were found for the effect of chemogenetic LC inhibition on decision making score (all ps≥0.115).

**Figure 3:**
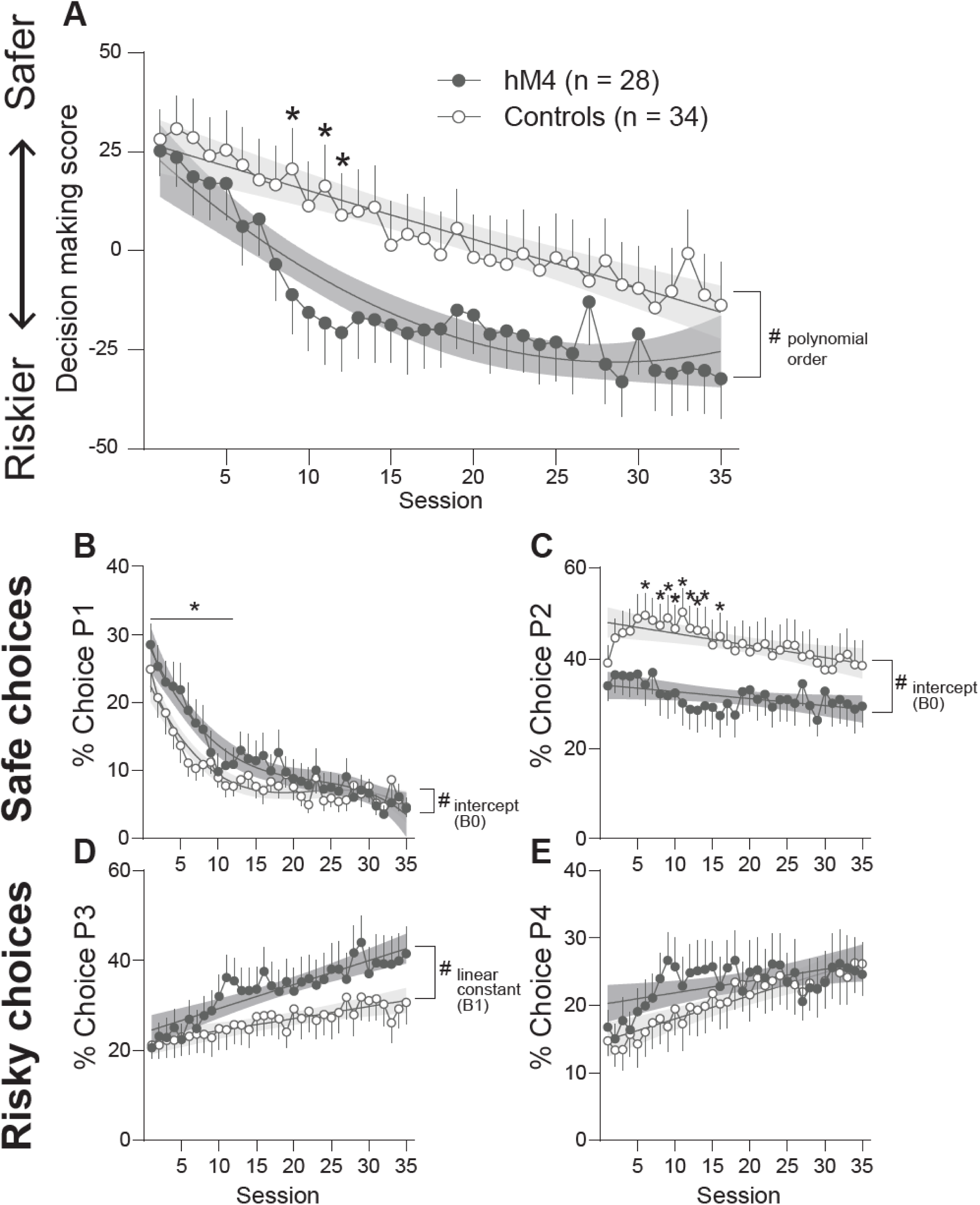
LC inhibition during crGT acquisition promotes risk-taking on the crGT. **A)** Chemogenetic inhibition of catecholaminergic LC cells impaired decision making on the crGT as indicated by lower decision making scores in the hM4 group during select early acquisition sessions. Decision making score also dropped more rapidly during early training in the hM4 group compared to controls, which instead exhibited a moderate and linear decline in score across acquisition. **B)** hM4 rats showed greater preference for the safest option, P1, during the early block of acquisition, and had an initially higher P1 preference as indicated by a greater intercept of the equation of best fit. **C)** Rats in the hM4 group failed to develop a preference for the most lucrative option, P2, during early to mid crGT acquisition. The line of best fit for P2 preference for controls was also significantly more elevated than that for hM4 rats. **D)** While preference for the risky option P3 did not differ between groups on any given session, hM4 rats exhibited a steeper increase in risky P3 choice across acquisition compared to controls. **E)** Chemogenetic inhibition of the LC did not influence choice of the riskiest option P4 at the level of the cohort. * p<0.05 hM4 vs controls on the indicated session according to a planned comparison. * and line: p<0.05 effect of group for indicated block of training. # p<0.05 group difference for at least one parameter for the equation of best fit – the parameter(s) that differed is (are) indicated on the figure. All data are presented as mean ± SEM and regression line or curve of best fit ± 95% confidence interval.

The greater decline in score observed in the hM4 group was accompanied by differences in preference for individual choice options between hM4 and control groups (choice x block x session x group: F_66,3036_=1.469, p=0.041, ηp^2^=0.031). Rats in the hM4 group differed in choice of the least risky option, choosing P1 more than controls throughout early and middle blocks of training (Figure 3B; block x group: F_2,92_=3.461, p=0.043, ηp^2^=0.070; group- early block: F_1,46_=5.627, p=0.022, ηp^2^=0.109; middle block: F_1,46_=7.369, p=0.006, ηp^2^=0.138; late block: F_1,46_=1.365, p=0.249). A cubic function best fit the trajectory of P1 choice for both control and hM4 groups (Figure 3B; block x session x group- linear contrast: F_1,46_=3.709, p=0.060, ηp^2^=0.075; cubic contrast: F_1,46_=5.567, p=0.023, ηp^2^=0.108; linear fit- hM4: F_1,976_=12.88, p<0.001; controls: F_1,1186_=29.63, p<0.001), yet the curves of best fit differed (same curve: F_4,2162_=8.59, p<0.001) such that P1 preference in the hM4 group was initially higher than that of controls (B0: F_1,2162_=4.937, p=0.026), despite a similar trend of P1 decline across sessions (all other coefficients: Fs≤0.048, ps≥0.827).

The hM4 subjects also selected the most lucrative option P2 significantly less than controls throughout training sessions 6 through 16 (Figure 3C; block x session x group: F_22,1012_=2.271, p=0.024, ηp^2^=0.047; hM4 vs controls: all ps≤0.044, ds≥0.524), excluding sessions 7 and 15 which failed to reach significance at the level of the post hoc comparison (hM4 vs controls: all ps≥0.065, ds≤0.479). The trajectory of P2 choice across training also differed – while data from both groups followed linear functions with comparable slopes (Figure 3C; block x session x group- linear contrast: F_1,46_=5.176, p=0.028, ηp^2^=0.101; same curve: F_2,2166_=51.48, p<0.001; B1: −0.155 (hM4) vs −0.269 (control), F_1,2166_=0.923, p=0.337), P2 preference in the control group was higher overall than P2 choice of the hM4 groups (B0: 34.26 (hM4) vs 48.38 (control), F_1,2166_=33.42, p<0.001).

The hM4 and control groups did not differ in choice of the risky options P3 or P4 on any specific session (Figure 3D,E; all group-wise interactions: Fs≤1.130, ps≥0.335), yet hM4 rats gained preference for P3 at a steeper rate than controls (Figure 3D; block x session x group linear contrast: F_1,46_=4.479, p=0.039, ηp^2^=0.076; same line: F_2,2166_=22.92, p<0.001; B1: 0.533 (hM4) vs 0.269 (control), F_1,2166_=5.690, p=0.0172).

### LC inhibition increases risk taking by promoting strategy shifts following safe wins and persistence at risky options irrespective of prior outcome

Following safe wins, hM4 animals showed a greater tendency than controls to switch from safe to risky choices (Figure 4A; block x session x group: F_22,814_=1.691, p=0.025, ηp^2^=0.044; early block- session x group: F_11,462_=1.804, p=0.051, ηp^2^=0.041; all other blocks: Fs≤1.504, ps≥0.179). Following losses on safe options, hM4 and control rats exhibited comparable switching behaviour (Figure 4B; all Fs≤2.760, ps≥0.074). Compared to controls, hM4 rats were less likely to switch away from risky options following both risky wins (Figure 4C; block x session x group: F_22,726_=2.110, p=0.038, ηp^2^=0.060; early block- session x group: F_11,407_=3.664, p=0.003, ηp^2^=0.090; all other blocks: Fs≤1.969, ps≥0.072) and risky losses (Figure 4D; block x session x group: F_22,858_=2.335, p=0.015, ηp^2^=0.056; early block- session x group: F_11,462_=2.245, p=0.050, ηp^2^=0.051; all other blocks: Fs≤1.321, ps≥0.210).

**Figure 4:**
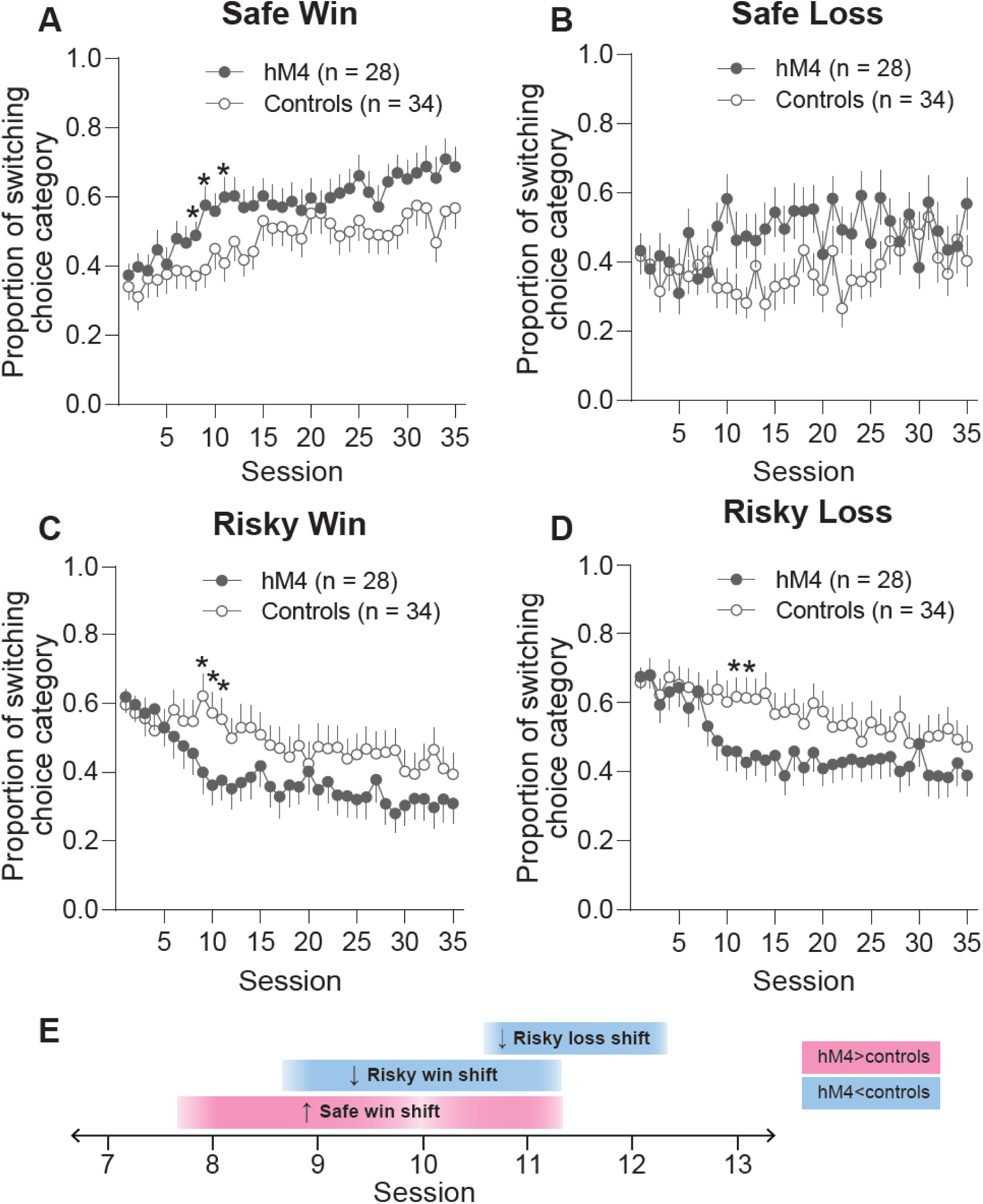
LC inhibition alters win-stay/lose-shift tendencies at varying timepoints during crGT acquisition. **A)** Rats in the hM4 group showed an increased tendency to shift choice strategy following safe wins on select sessions in early acquisition, but **B)** did not significantly influence shifting behaviour following losses from safe options. hM4 rats also exhibited a reduced tendency to shift away from risky choices following either **C)** risky wins and **D)** risky losses. **E)** These alterations in trial-by-trial behaviour emerged on distinct sessions during crGT acquisition. The greater safe win-shift behaviour of hM4 rats appeared first, on session 8, and overlapped with the reduced risky win-shift tendency, which appeared on the following session 9. The reduction in risky loss-shift behaviour occurred last on session 11. * p<0.05 hM4 vs controls on the indicated session according to a planned comparison. All data a presented as mean ± SEM.

Notably, differences in trial-by-trial behaviour between hM4 and control rats appeared transiently and emerged on distinct sessions across acquisition. Initially hM4 rats exhibited greater shifting tendencies following safe wins from session 8 (Figure 4E; hM4 vs controls-sessions 8, 9 & 11: ps≤0.050, ds≥0.516; all other sessions ps≥0.094, ds≤0.441), prior to the observed reduction in choice shifts following risky wins from session 9 (Figure 4E; hM4 vs control- sessions 9,10&11: ps≤0.038, ds≥0.541; all other sessions: ps≥0.078, ds≤0.457). Lastly, reduced shifting behaviour of hM4 rats following risky losses emerged on session 11 (Figure 4E; hM4 vs control- sessions 11&12: ps≤0.054, ds≥0.506; all other sessions: ps≥0.060, ds≤0.485).

No within-subject contrasts reached significance for trial-by-trial data (group-wise effects and interactions: all Fs≤3.501, ps≥0.069), and thus, trajectories of lose-shift behaviour across acquisition were not compared between groups.

### LC inhibition selectively enhances motor impulsivity in females

Chemogenetic inhibition of the LC selectively increased motor impulsive premature responding in female rats during the early block of crGT training (Figure 5A; block x sex x group: F_2,92_=3.568, p=0.044, ηp^2^=0.072; group- females early block: F_1,24_=6.303, p=0.019, ηp^2^=0.208; all other blocks all Fs≤0.572, ps≥0.475; males: all Fs≤1.070, ps≥0.410). As indicated by different lines of best fit (block x session x group x sex linear contrast: F_1,46_=4.252, p=0.045, ηp^2^=0.085; females- block x session x group linear contrast: F_1,24_=7.064, p=0.014, ηp^2^=0.227; same line: F_2,1116_=72.75, p<0.001), hM4 females demonstrate higher initial levels of premature responding (Figure 5A; B0: 38.22 (hM4) vs 22.00 (control), F_1,1116_=90.23, p<0.001) and a steeper decline across training (Figure 5A; B1: −0.502 (hM4) vs −0.118 (control), F_1,1116_=21.59, p<0.001) compared to control females. LC inhibition did not influence the trajectory of premature responding in males (Figure 5B; block x session x group linear contrast: F_1,24_=1.367, p=0.225).

**Figure 5:**
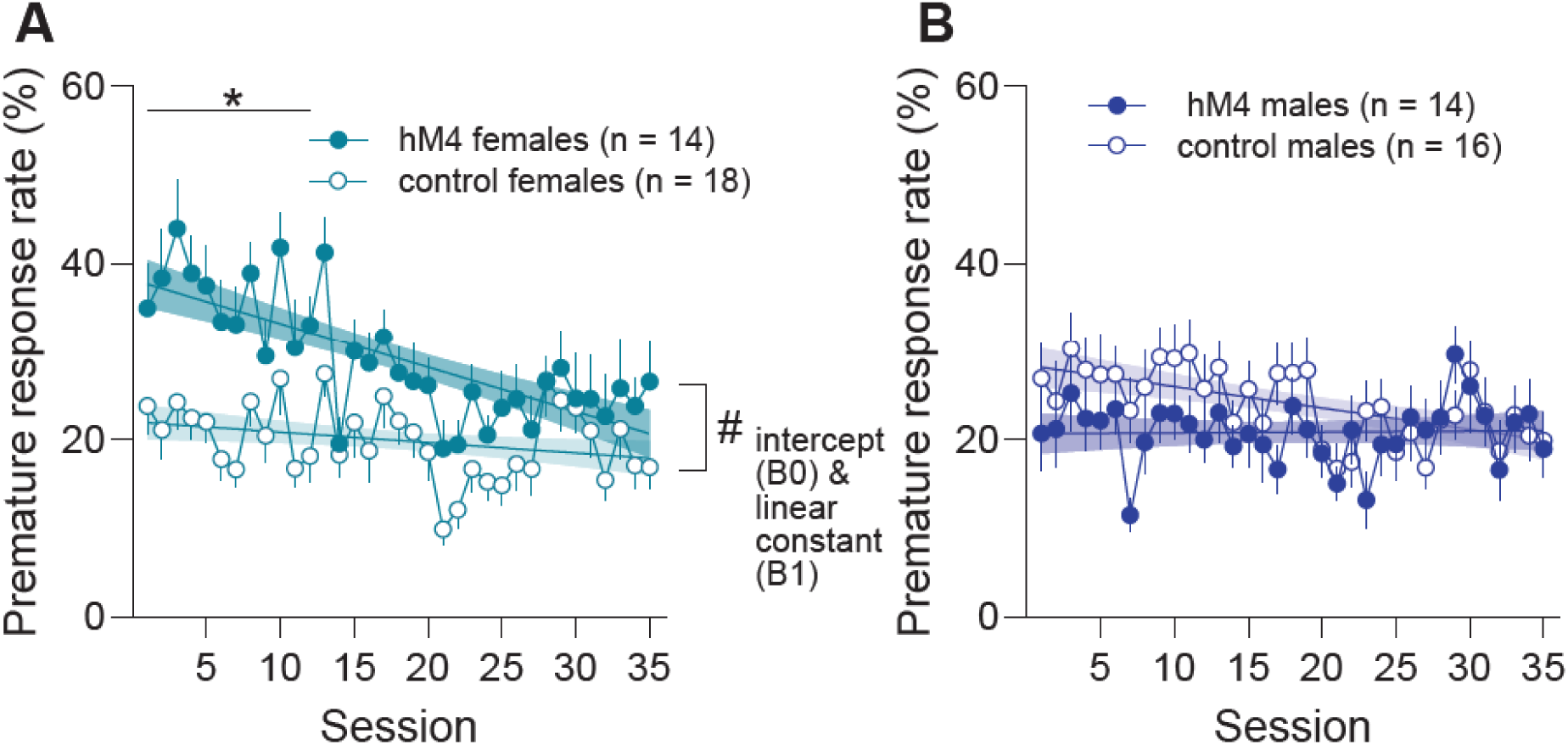
Chemogenetic inhibition of LC selectively increases motor impulsivity in females. **A)** Irrespective of impulsivity level, LC inhibition enhanced impulsive premature responding in females – a transient effect which was observed only during the early block of training. **B)** LC inhibition did not influence motor impulsivity in males. * and line: p<0.05 effect of group for indicated block of training. # p<0.05 group difference for at least one parameter for the equation of best fit – the parameter that differed is indicated on the figure. All data a presented as mean ± SEM and regression line or curve of best fit ± 95% confidence interval.

### LC inhibition has marginal effects on latencies and number of trials completed and omitted

LC inhibition increased the latency to collect food rewards (Figure 6A; group: F_1,46_=6.617, p=0.013, ηp^2^=0.126), but did not alter the latency to make a choice (Figure 6B; all group-wise effects: all Fs≤3.588, ps≥0.064) or the number of omitted trials (Figure 6C; group-wise effects: all Fs≤1.827, ps≥0.075). The number of completed trials was lower in hM4 rats during the early block of crGT acquisition (Figure 5D; block x group: F_2,92_=5.158, p=0.012, ηp^2^=0.038; group- early block: F_1,46_=5.761, p=0.020, ηp^2^=0.111; middle and late blocks: all Fs≤1.041, ps≥0.313), consistent with the longer latency to collect food rewards (Figure 6A) and greater preference for risky options which yield longer timeout punishments in the hM4 group (Figure 3A).

**Figure 6:**
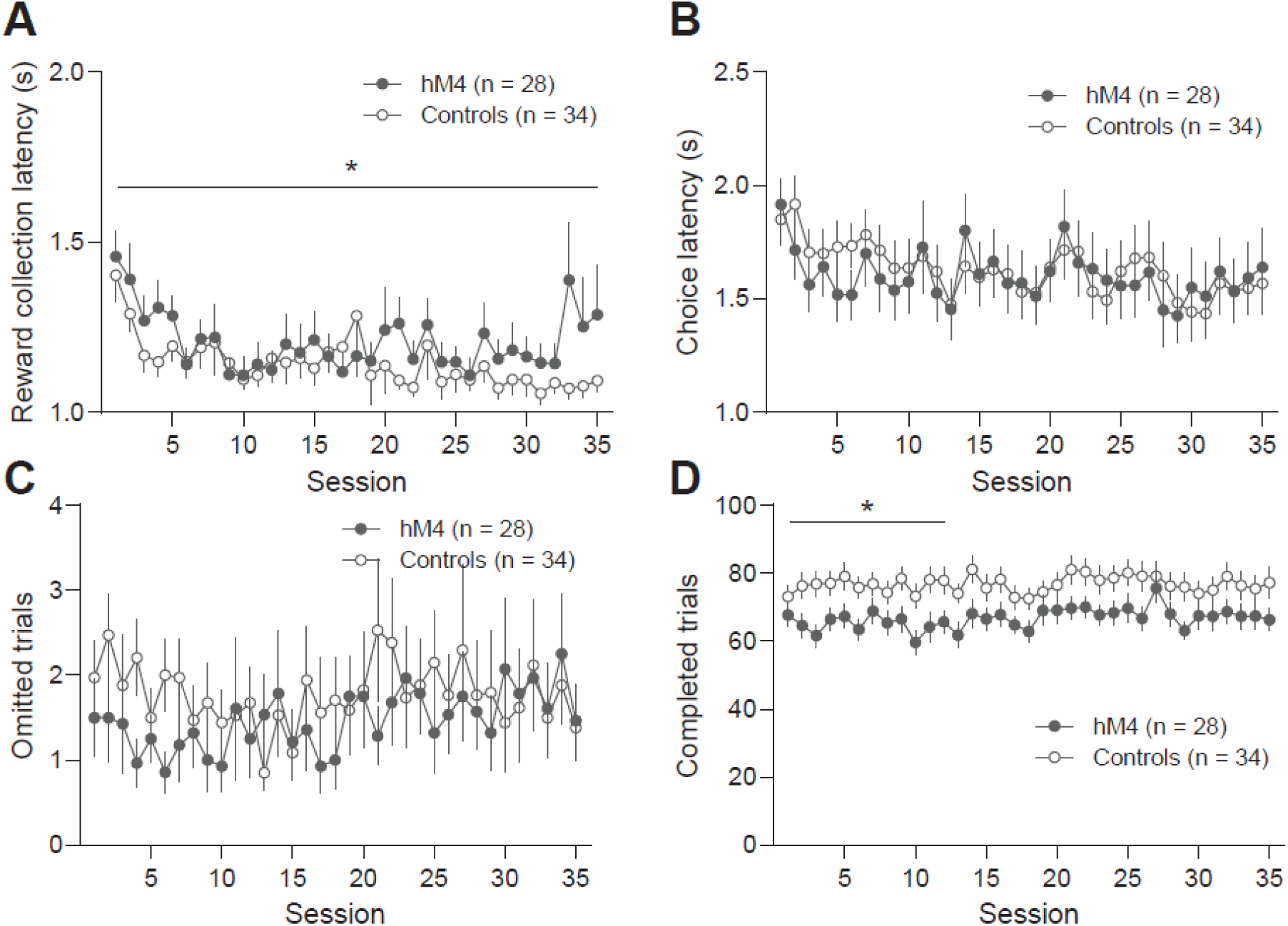
Inhibition of the LC reduces speed of reward collection and the number of completed trials. **A)** hM4 rats take longer than controls to collect food reward – an effect which persists throughout crGT training. Chemogenetic LC inhibition did not influence **B)** the latency to make a choice nor **C)** the number of trials omitted due to failure to make a choice. **D)** LC inhibition decreased the number of completed trials, consistent with the hM4 group’s increased collection latency and preference for high-risk options which incur longer timeout punishments. * and line: p<0.05 effect of group for indicated block of training. All data a presented as mean ± SEM.

### Behavioural effect of LC inhibition on collection latency washes out

By the end of acquisition, crGT performance between hM4 and control groups differed only by collection latency (see above results section). By session six of post-washout CNO-free testing, crGT behaviour reached statistical stability for all behavioural variables across five consecutive sessions (session: all Fs≤3.144, ps≥0.081). Performance of hM4 and control groups during these CNO-free sessions did not differ for any variable (Supp Figure 1A-J; all group-wise effects: all Fs≤2.118, ps≥0.094).

## Discussion

Here we show that chemogenetic suppression of putative noradrenergic LC cells impaired performance on the crGT, irrespective of sex, by hastening the rate at which rats develop risky strategies across task acquisition. This was driven by a failure to adopt a preference for the most lucrative option, P2, and a faster increase in preference for risky option P3. Suppression of putative noradrenergic LC neurons promoted sampling of risky options following a safe win and reduced the likelihood of switching away from risky options, even if a loss occurred. Differences in win-shift behaviour emerged prior to altered lose-shift tendencies, indicating that LC inhibition first impacts the way in which wins influence choice before altering behavioural sensitivity to losses. Females showed exaggerated impulsive premature responding following LC inhibition – an effect which was not observed in males. LC suppression also rendered all animals somewhat slower to collect food pellet rewards.

The LC fundamentally regulates sensory processing (Waterhouse & Navarra, 2019), a critical function to consider during performance on a task that activates multiple sensory modalities (i.e., gustation and olfaction via food rewards; salient audiovisual stimuli). However, hM4-mediated inhibition of the LC leaves auditory startle responses (Kabanova et al., 2024) and visual processing speed intact (Fitzpatrick et al., 2019). Further, win-paired cues promote risk preference (Barrus & Winstanley, 2016). If LC inhibition had impaired sensory processing of these cues, decision making should arguably have become more optimal, resembling that seen on the uncued rGT, and opposite to the increase in risk-taking observed here.

Noradrenergic neurons within the LC also regulate general arousal. If chemogenetic inhibition of this neuronal population had caused arousal to decline substantially, we would have expected omissions and response latencies to increase, while trials completed and premature responding declined. In contrast, we observed a selective increase in reward collection latency, and a temporary elevation in premature responding in females. While we did observe a reduction in trials with our experimental manipulation, this can be better explained by the increased selection of risky options during the same time period, rather than any wholesale changes in arousal levels.

Such null effects on sensory processing or arousal- canonical functions of the LC-following chemogenetic inhibition of noradrenergic LC neurons may call into question the efficacy of this manipulation. The effects of LC inhibition on both decision making and impulsivity were only detected during relatively early training sessions, which may indicate that sufficient functional compensation had occurred by later sessions as to negate the effects of this manipulation. However, reward collection latency was generally higher in hM4-expressing animals throughout the CNO dosing period, and only returned to control levels when CNO dosing ceased, indicating chemogenetic inhibition of LC neurons remained in effect for the duration of the experiment. The temporal specificity of the effects on choice and impulse control may instead reflect a more prominent role for LC activity in regulating these functions when animals are first learning to adapt their behaviour to the task demands.

It is important to note that chemogenetic suppression of LC neurons does not completely silence the LC (Markovic et al., 2024; Vazquez et al., 2025). As such, firing should be less probable to weak stimuli, but still achievable with strong excitatory input (Smith et al., 2016). The LC facilitates reward learning through encoding reward prediction errors (RPEs) (Breton-Provencher et al., 2022; Brink et al., 2025; Su & Cohen, 2022) in situations where reinforcement schedules are lean and outcomes are predictable (Breton-Provencher et al., 2022), and may therefore serve as a node that shapes and titrates conditioned responses to motivationally-relevant cues that are situationally rare (Bouret & Richmond, 2015; Bouret & Sara, 2004). LC responses to probabilistic outcomes scale inversely with likelihood, such that less probable outcomes elicit larger LC RPEs (Breton-Provencher et al., 2022; Jordan & Keller, 2023). As such, unlikely risky wins should be accompanied by more robust LC activation as compared to the more probable safe wins in the crGT. Chemogenetic inhibition may effectively block weaker LC activation to more certain rewards (ie. from P1/P2), while failing to prevent stronger LC responses to unlikely risky wins, thereby disproportionately enhancing reinforcement learning following risky wins. Likewise, hM4-mediated LC inhibition may completely attenuate smaller RPE signals arising from frequent risky losses, while stronger loss signals following rare safe losses may persist, emphasizing punishment learning from safe losses. The blunted LC encoding of risky losses, alongside relatively preserved loss signals on safe losses, could simultaneously explain the enhanced risky P3 preference and reduced optimal P2 choice observed when the LC is inhibited, respectively.

This theoretical framework whereby LC inhibition reduces the impact of likely outcomes (ie. safe wins and risky losses) on decision making is also generally consistent with our trial-by-trial data. Our WSLS analyses indicate that LC-inhibition promotes persistence at risky options, where losses are likely, and shifts choice away from safe options which yield much more probable wins, consistent with underrepresentation of likely outcomes in the decision making landscape. The WSLS data are also consistent with the suggested ‘unsigned’ nature of LC RPEs (Jordan, 2024; Jordan & Keller, 2023; Sales et al., 2019), given that LC inhibition did not render behaviour preferentially sensitive to either wins or losses. Additionally, the WSLS data indicate that LC inhibition does not elicit an overall reduction in behavioural flexibility, as would be evidenced by decreased shift likelihood following any outcome. The lack of effect on safe losses may be partially driven by the large variance in safe loss trial-by-trial data, given the relatively low probability of encountering a safe loss during a crGT session.

The presence of salient win-paired cues themselves promotes risk-taking on the crGT (Barrus & Winstanley, 2016; Winstanley & Hynes, 2021), and perhaps the LC contributes to this behavioural phenomenon. Indeed, the risk promoting effect of these cues is modulated by noradrenergic drugs (Baarendse et al., 2014; Chernoff et al., 2021; Silveira et al., 2016), consistent with the current data. Strong phasic LC responses to less frequent but highly salient task-relevant cues (Grant et al., 1988; Rajkowski et al., 2004; Vazey et al., 2018) may not be completely silenced by hM4-mediated inhibition (Smith et al., 2016), and thus may still contribute to cue-invigorated risk-taking in the experimental group. Phasic LC activity is associated with selective attention and behavioural rigidity (Aston-Jones et al., 1999; Aston-Jones & Cohen, 2005), and may contribute to persistent choice of highly-cued risky options on the crGT in part through these processes. The observed increase in reward collection latency following LC inhibition loosely supports this interpretation, since enhanced attention to the salient win-concurrent cues on the wall opposite to the food magazine could contribute to the elongated reward collection latency. Further, it is unlikely that chemogenetic LC suppression increased collection latency here by altering naturalistic feeding behaviour, given that LC *activation* suppresses feeding, yet food intake remains intact following hM4-mediated LC-NA inhibition (Sciolino et al., 2022).

In females, we highlight the particular importance of LC-NA for impulse control, as evidenced by a female-selective exacerbation of motor impulsivity following LC inhibition. This is consistent with pharmacological crGT data, whereby compounds and doses that significantly altered impulsive premature responding in females were without effect on this variable in males (Chernoff et al., 2021). In contrast, males were more sensitive to the anti-impulsivity effect of noradrenaline transporter blockade in an attentional reaction time task (Mei et al., 2021). Sex differences in behavioural sensitivity appears to depend on the nature of the task and the associated attentional demands.

Numerous reports indicate sex differences in LC morphology and responsivity to stress which may render the female LC more excitable (Bangasser et al., 2019; Curtis et al., 2006; Mariscal Ramírez et al., 2023; Pinos et al., 2001). Enhanced LC burst firing is associated with greater motor impulsivity (Cieślak et al., 2017). If hM4-mediated LC suppression were to less effectively blunt the stronger LC activation of females compared to males, while effectively reducing noradrenergic tone in both sexes, this could explain why we only see enhanced impulsivity in inhibited females. The relationship between LC-NA activity and impulsivity is likely not linear, as remarkably low or high noradrenaline levels may similarly impair top-down impulse control (Aston-Jones & Cohen, 2005; Chamberlain & Robbins, 2013). To our knowledge, no study has directly demonstrated an inverted-U relationship between LC-NA activity and impulse control, but it would be valuable to document the impact of chemogenetic excitation of LC-NA on this behaviour.

Similar sex differences have been reported in dopaminergic modulation of motor impulsivity, whereby chemogenetic inhibition and excitation of midbrain dopamine influences impulsivity selectively in males or females, respectively (Hynes et al., 2021, 2024). Chemogenetic stimulation of midbrain dopamine neurons increased impulsivity in females similarly to chemogenetic LC inhibition in the present study. This could indicate an antagonistic but complementary contribution of dopamine and noradrenaline to impulse control in females. LC-NA can indirectly influence the mesolimbic dopamine system (Aono et al., 2015; Mitrano et al., 2012; Park et al., 2017), but it remains to be determined whether noradrenergic modulation of dopaminergic activity is responsible for the increases in impulsivity observed here.

With its extensive efferent projections, the LC could influence performance of the crGT through myriad circuits. The LC may guide decision-making and impulse control through direct action in frontal regions such as the orbitofrontal cortex and medial prefrontal cortex, which mediate the effects of various noradrenergic compounds on choice and impulsivity on the crGT (Chernoff et al., 2024). Alternatively, the LC projects selectively to the nucleus accumbens shell within the striatum, a key mediator of noradrenaline’s influence over motor impulsivity (Economidou et al., 2012; Velazquez-Sanchez et al., 2023). Determining the circuit-level mechanisms through which LC-NA acts to orchestrate risk-taking remains an important goal.

In sum, we provide causal evidence that the noradrenergic LC is required for the acquisition of optimal decision-making strategies using a translationally-relevant model of gambling-like behaviour, such that impaired LC function promotes maladaptive risk taking during early task acquisition. Such choice patterns parallel the impaired decision making observed across numerous human psychiatric conditions such as substance use disorder (SUD), major depression, and gambling disorder, and suggest dysregulated LC function could contribute to these decision-making deficits. We also show that the LC regulates impulse control selectively in females, which may contribute to sex-dependent variation across psychiatric disorders in which impulsivity is a prominent feature (ex. SUD, eating disorders). Our findings provide novel insights into the contribution of the LC-NA system to complex behaviour, and underscore the functional differences in LC regulation of behaviour depending on biological sex.

## Acknowledgements and funding information

This work was supported by an NSERC Discovery Grant awarded to CAW (RGPIN-2023-04030). CSC was supported by a Canadian Graduate Scholarship- Master’s level and TJH was supported by a Marshall Graduate Scholarship. This work took place at a UBC campus situated on the traditional, ancestral, and unceded land of the x&#x25A1;məθk&#x25A1;əy&#x25A1; əm (Musqueam), sə&#x25A1;lílwəta&#x25A1;&#x25A1;Selilwitulh (Tsleil-Waututh) and S&#x25A1;wx&#x25A1;wú7mesh (Squamish) Peoples. We acknowledge and are grateful for their stewardship of this land for thousands of years

## Data availability statement

Data are available upon request. Please contact the corresponding author CSC with requests.

## Supplemental Figure legends

*Supplemental* figure 1*: Behavioural effects of chemogenetic LC inhibition wash out effectively.* Following a 7-day period of washout (indicated by grey), hM4 and control rats exhibit comparable performance on **A-E)** decision making, **F)** impulsivity, **G-H)** latency, and **I-J)** trial variables when tested CNO-free on the crGT. All data a presented as mean ± SEM.

*Supplemental* figure 2*: TG+ and TG- controls exhibit comparable behaviour on the crGT.* **A-J)** Cre-dependent expression of a mCherry control fluorophore in the LC did not influence behaviour on the crGT as transgene positive (TG+) and negative (TG-) TH::Cre rats show comparable behaviour on all crGT variables throughout acquisition, and thus were pooled as controls. All data a presented as mean ± SEM.

## References

Ansquer, S., Belin-Rauscent, A., Dugast, E., Duran, T., Benatru, I., Mar, A. C., Houeto, J. L., & Belin, D. (2014). Atomoxetine decreases vulnerability to develop compulsivity in high impulsive rats. Biological Psychiatry, 75(10), 825–832. 10.1016/j.biopsych.2013.09.031

Aono, Y., Taguchi, H., Saigusa, T., Uchida, T., Takada, K., Takiguchi, H., Shirakawa, T., Shimizu, N., Koshikawa, N., & Cools, A. R. (2015). Simultaneous activation of the α1A-, α1B- and α1D-adrenoceptor subtypes in the nucleus accumbens reduces accumbal dopamine efflux in freely moving rats. Behavioural Pharmacology, 26(1 and 2-Special Issue), 73. 10.1097/FBP.0000000000000113

Aston-Jones, G., & Bloom, F. E. (1981). Nonrepinephrine-containing locus coeruleus neurons in behaving rats exhibit pronounced responses to non-noxious environmental stimuli. Journal of Neuroscience, 1(8), 887–900.

Aston-Jones, G., & Cohen, J. D. (2005). An integrative theory of locus coeruleus-norepinephrine function: Adaptive gain and optimal performance. Annual Reviews of Neuroscience, 28, 403–450. 10.1146/annurev.neuro.28.061604.135709

Aston-Jones, G., Rajkowski, J., & Cohen, J. (1999). Role of locus coeruleus in attention and behavioral flexibility. Biological Psychiatry, 46(9), 1309–1320. 10.1016/S0006-3223(99)00140-7

Aston-Jones, G., Rajkowski, J., Kubiak, P., & Alexinsky, T. (1994). Locus coeruleus neurons in monkey are selectively activated by attended cues in a vigilance task. Journal of Neuroscience, 14(7), 4467–4480.

Baarendse, P. J. J., Winstanley, C. A., & Vanderschuren, L. J. M. J. (2014). Simultaneous blockade of dopamine and noradrenaline reuptake promotes disadvantageous decision making in a rat gambling task. Psychopharmacology, 225(3), 719–731. 10.1007/s00213-012-2857-z.Simultaneous

Bangasser, D. A., Curtis, A., Reyes, B. A. S., Bethea, T. T., Parastatidis, I., & Ischiropoulos, H. (2010). Sex differences in corticotropin-releasing factor receptor signaling and trafficking: Potential role in female vulnerability to stress-related psychopathology.\ Molecular Psychiatry, 15(9), 896–904. 10.1038/mp.2010.66

Bangasser, D. A., Eck, S. R., & Ordoñes Sanchez, E. (2019). Sex differences in stress reactivity in arousal and attention systems. Neuropsychopharmacology, 44(1), Article 1. 10.1038/s41386-018-0137-2

Bangasser, D. A., Reyes, B. A. S., Piel, D., Garachh, V., Zhang, X. Y., Plona, Z. M., Van Bockstaele, E. J., Beck, S. G., & Valentino, R. J. (2013). Increased vulnerability of the brain norepinephrine system of females to corticotropin-releasing factor overexpression. Molecular Psychiatry, 18(2), 166–173. 10.1038/mp.2012.24

Bangasser, D. A., Zhang, X., Garachh, V., Hanhauser, E., & Valentino, R. J. (2011). Sexual dimorphism in locus coeruleus dendritic morphology: A structural basis for sex differences in emotional arousal. Physiology & Behavior, 103(3–4), 342–351. 10.1016/j.physbeh.2011.02.037

Barrus, M. M., & Winstanley, C. A. (2016). Dopamine D3 receptors modulate the ability of win-paired cues to increase risky choice in a rat gambling task. Journal of Neuroscience, 36(3), 785–794. 10.1523/JNEUROSCI.2225-15.2016

Benn, A., & Robinson, E. S. J. (2017). Differential roles for cortical versus sub-cortical noradrenaline and modulation of impulsivity in the rat. Psychopharmacology, 234(2), 255–266. 10.1007/s00213-016-4458-8

Bouret, S., & Richmond, B. J. (2015). Sensitivity of Locus Ceruleus Neurons to Reward Value for Goal-Directed Actions. Journal of Neuroscience, 35(9), 4005–4014. 10.1523/JNEUROSCI.4553-14.2015

Bouret, S., & Sara, S. J. (2004). Reward expectation, orientation of attention and locus coeruleus-medial frontal cortex interplay during learning. European Journal of Neuroscience, 20(3), 791–802. 10.1111/j.1460-9568.2004.03526.x

Breton-Provencher, V., Drummond, G. T., Feng, J., Li, Y., & Sur, M. (2022). Spatiotemporal dynamics of noradrenaline during learned behaviour. Nature, 606(7915), 732–738. 10.1038/s41586-022-04782-2

Brink, A. K., Lui, S. K. C., & Corbit, L. H. (2025). Alpha-2 agonism in the locus coeruleus impairs learning driven by negative prediction error. Neuropsychopharmacology, 50(7), 1186–1193. 10.1038/s41386-025-02092-5

Carvalheiro, J., & Philiastides, M. G. (2023). Distinct spatiotemporal brainstem pathways of outcome valence during reward- and punishment-based learning. Cell Reports, 42(12). 10.1016/j.celrep.2023.113589

Chamberlain, S. R., & Robbins, T. W. (2013). Noradrenergic modulation of cognition: Therapeutic implications. Journal of Psychopharmacology, 27(8), 694–718. 10.1177/0269881113480988

Chernoff, C. S., Belin-Rauscent, A., Puaud, M., Torrisi, S. A., Fouyssac, M., Németh, B., Yu, C. Z., Higuera-Matas, A., Jones, S., & Belin, D. (2025). Compulsive coping behaviour, developed predominantly by sign-trackers, is exacerbated by chronic atomoxetine. Biological Psychiatry, 0(0). 10.1016/j.biopsych.2025.07.012

Chernoff, C. S., Hynes, T. J., Schumacher, J. D., Ramaiah, S., Avramidis, D. K., Mortazavi, L., Floresco, S. B., & Winstanley, C. A. (2024). Noradrenergic regulation of cue-guided decision making and impulsivity is doubly dissociable across frontal brain regions. Psychopharmacology, 241(4), 767–783. 10.1007/s00213-023-06508-2

Chernoff, C. S., Hynes, T. J., & Winstanley, C. A. (2021). Noradrenergic contributions to cue-driven risk-taking and impulsivity. Psychopharmacology, 238(7), 1765–1779. 10.1007/s00213-021-05806-x

Cieślak, P. E., Llamosas, N., Kos, T., Ugedo, L., Jastrzębska, K., Torrecilla, M., & Rodriguez Parkitna, J. (2017). The role of NMDA receptor-dependent activity of noradrenergic neurons in attention, impulsivity and exploratory behaviors. *Genes*, Brain and Behavior, 16(8), 812–822. 10.1111/gbb.12383

Curtis, A. L., Bethea, T., & Valentino, R. J. (2006). Sexually Dimorphic Responses of the Brain Norepinephrine System to Stress and Corticotropin-Releasing Factor. Neuropsychopharmacology, 31(3), 544–554. 10.1038/sj.npp.1300875

Economidou, D., Theobald, D. E. H., Robbins, T. W., Everitt, B. J., & Dalley, J. W. (2012). Norepinephrine and dopamine modulate impulsivity on the five-choice serial reaction time task through opponent actions in the shell and core sub-regions of the nucleus accumbens. Neuropsychopharmacology, 37(9), 2057–2066. 10.1038/npp.2012.53

Engberg, G., Oreland, L., Thorén, P., & Svensson, T. (1987). Locus coeruleus neurons show reduced alpha2-receptor responsiveness and decreased basal activity in spontaneously hypertensive rats. Journal of Neural Transmission, 69(1), 71–83. 10.1007/BF01244098

Fitzpatrick, C. M., Runegaard, A. H., Christiansen, S. H., Hansen, N. W., Jørgensen, S. H., McGirr, J. C., de Diego Ajenjo, A., Sørensen, A. T., Perrier, J. F., Petersen, A., Gether, U., Woldbye, D. P. D., & Andreasen, J. T. (2019). Differential effects of chemogenetic inhibition of dopamine and norepinephrine neurons in the mouse 5-choice serial reaction time task. Progress in Neuro-Psychopharmacology and Biological Psychiatry, 90, 264– 276. 10.1016/j.pnpbp.2018.12.004

Grant, S. J., Aston-Jones, G., & Redmond, D. E. (1988). Responses of primate locus coeruleus neurons to simple and complex sensory stimuli. Brain Research Bulletin, 21(3), 401–410. 10.1016/0361-9230(88)90152-9

Hathaway, B. A., Li, A., Brodie, H. G., Silveira, M. M., Tremblay, M., Seo, Y. S., & Winstanley, C. A. (2024). Dopamine activity in the nigrostriatal pathway alters cue-induced risky choice patterns in female rats. European Journal of Neuroscience, 59(7), 1621–1637. 10.1111/ejn.16287

Hesse, S., Müller, U., Rullmann, M., Luthardt, J., Bresch, A., Becker, G. A., Zientek, F., Patt, M., Meyer, P. M., Blüher, M., Strauß, M., Fenske, W., Hankir, M., Ding, Y. S., Hilbert, A., & Sabri, O. (2017). The association between in vivo central noradrenaline transporter availability and trait impulsivity. Psychiatry Research - Neuroimaging, 267, 9–14. 10.1016/j.pscychresns.2017.06.013

Hynes, T. J., Chernoff, C. S., Hrelja, K. M., Tse, M. T. L., Avramidis, D. K., Lysenko-Martin, M. R., Calderhead, L., Kaur, S., Floresco, S. B., & Winstanley, C. A. (2024). Win-Paired Cues Modulate the Effect of Dopamine Neuron Sensitization on Decision Making and Cocaine Self-administration: Divergent Effects Across Sex. *Biological Psychiatry*, *Reward Learning*, Circuits, and Addiction Risk, 95(3), 220–230. 10.1016/j.biopsych.2023.08.021

Hynes, T. J., Hrelja, K. M., Hathaway, B. A., Hounjet, C. D., Chernoff, C. S., Ebsary, S. A., Betts, G. D., Russell, B., Ma, L., Kaur, S., & Winstanley, C. A. (2021). Dopamine neurons gate the intersection of cocaine use, decision making, and impulsivity. Addiction Biology, 26(6), e13022. 10.1111/adb.13022

Jordan, R. (2024). The locus coeruleus as a global model failure system. Trends in Neurosciences, 47(2), 92–105. 10.1016/j.tins.2023.11.006

Jordan, R., & Keller, G. B. (2023). The locus coeruleus broadcasts prediction errors across the cortex to promote sensorimotor plasticity. eLife, 12, RP85111. 10.7554/eLife.85111

Kabanova, A., Yang, M., Logothetis, N. K., & Eschenko, O. (2024). Partial chemogenetic inhibition of the locus coeruleus due to heterogeneous transduction of noradrenergic neurons preserved auditory salience processing in wild-type rats. European Journal of Neuroscience, 60(9). 10.1111/ejn.16550

Mariscal, P., Bravo, L., Llorca-Torralba, M., Razquin, J., Miguelez, C., Suárez-Pereira, I., & Berrocoso, E. (2023). Sexual differences in locus coeruleus neurons and related behavior in C57BL/6J mice. Biology of Sex Differences, 14(1), 64. 10.1186/s13293-023-00550-7

Mariscal Ramírez, P., Bravo García, L., Llorca Torralba, M., Razquin, J., Miguelez, C., Suárez-Pereira, I., & Berrocoso Domínguez, E. M. (2023). Sexual differences in locus coeruleus neurons and related behavior in C57BL/6J mice. Biology of Sex Differences, *Vol.* 14, Núm. 1, 2023. 10.1186/S13293-023-00550-7

Markovic, T., Higginbotham, J., Ruyle, B., Massaly, N., Yoon, H. J., Kuo, C.-C., Kim, J. R., Yi, J., Garcia, J. J., Sze, E., Abt, J., Teich, R. H., Dearman, J. J., McCall, J. G., & Morón, J. A. (2024). A locus coeruleus to dorsal hippocampus pathway mediates cue-induced reinstatement of opioid self-administration in male and female rats. Neuropsychopharmacology, 1–9. 10.1038/s41386-024-01828-z

Mei, X., Wang, L., Yang, B., & Li, X. (2021). Sex differences in noradrenergic modulation of attention and impulsivity in rats. Psychopharmacology, 238(8), 2167–2177. 10.1007/s00213-021-05841-8

Mitrano, D. A., Schroeder, J. P., Smith, Y., Cortright, J. J., Bubula, N., Vezina, P., & Weinshenker, D. (2012). Alpha-1 Adrenergic Receptors are Localized on Presynaptic Elements in the Nucleus Accumbens and Regulate Mesolimbic Dopamine Transmission. Neuropsychopharmacology, 37(9), 2161–2172. 10.1038/npp.2012.68

Mortazavi, L., Hynes, T. J., Chernoff, C. S., Ramaiah, S., Brodie, H. G., Russell, B., Hathaway, B. A., Kaur, S., & Winstanley, C. A. (2023). D2/3 Agonist during Learning Potentiates Cued Risky Choice. Journal of Neuroscience. 10.1523/JNEUROSCI.1459-22.2022

Mulvey, B., Bhatti, D. L., Gyawali, S., Lake, A. M., Kriaucionis, S., Ford, C. P., Bruchas, M. R., Heintz, N., & Dougherty, J. D. (2018). Molecular and Functional Sex Differences of Noradrenergic Neurons in the Mouse Locus Coeruleus. Cell Reports, 23(8), 2225–2235. 10.1016/j.celrep.2018.04.054

Navarra, R., Graf, R., Huang, Y., Logue, S., Comery, T., Hughes, Z., & Day, M. (2008). Effects of atomoxetine and methylphenidate on attention and impulsivity in the 5-choice serial reaction time test. Progress in Neuro-Psychopharmacology and Biological Psychiatry, 32(1), 34–41. 10.1016/j.pnpbp.2007.06.017

Park, J. W., Bhimani, R. V., & Park, J. (2017). Noradrenergic Modulation of Dopamine Transmission Evoked by Electrical Stimulation of the Locus Coeruleus in the Rat Brain. ACS Chemical Neuroscience, 8(9), 1913–1924. 10.1021/acschemneuro.7b00078

Pinos, H., Collado, P., Rodrı guez-Zafra, M., Rodrı guez, C., Segovia, S., & Guillamón, A. (2001). The development of sex differences in the locus coeruleus of the rat. Brain Research Bulletin, 56(1), 73–78. 10.1016/S0361-9230(01)00540-8

Rajkowski, J., Majczynski, H., Clayton, E., & Aston-Jones, G. (2004). Activation of monkey locus coeruleus neurons varies with difficulty and performance in a target detection task. Journal of Neurophysiology, 92(1), 361–371. 10.1152/jn.00673.2003

Robinson, E. S. J., Eagle, D. M., Mar, A. C., Bari, A., Banerjee, G., Jiang, X., Dalley, J. W., & Robbins, T. W. (2008). Similar effects of the selective noradrenaline reuptake inhibitor atomoxetine on three distinct forms of impulsivity in the rat. Neuropsychopharmacology, 33(5), 1028–1037. 10.1038/sj.npp.1301487

Russell, V. A., Allie, S., & Wiggins, T. (2000). Increased noradrenergic activity in prefrontal cortex slices of an animal model for attention-deficit hyperactivity disorder—The spontaneously hypertensive rat. Behavioural Brain Research, 117(1), 69–74. 10.1016/S0166-4328(00)00291-6

Sales, A. C., Friston, K. J., Jones, M. W., Pickering, A. E., & Moran, R. J. (2019). Locus Coeruleus tracking of prediction errors optimises cognitive flexibility: An Active Inference model. PLoS Computational Biology, 15(1), e1006267.

Sara, S. J., & Bouret, S. (2012). Orienting and Reorienting: The Locus Coeruleus Mediates Cognition through Arousal. Neuron, 76(1), 130–141. 10.1016/j.neuron.2012.09.011

Sciolino, N. R., Hsiang, M., Mazzone, C. M., Wilson, L. R., Plummer, N. W., Amin, J., Smith, K. G., McGee, C. A., Fry, S. A., Yang, C. X., Powell, J. M., Bruchas, M. R., Kravitz, A. V., Cushman, J. D., Krashes, M. J., Cui, G., & Jensen, P. (2022). Natural locus coeruleus dynamics during feeding. Science Advances, 8(33), eabn9134. 10.1126/sciadv.abn9134

Selden, N., Cole, B. J., Everitt, B. J., & Robbins, T. W. (1990). Damage to ceruleo-cortical noradrenergic projections impairs locally cued but enhances spatially cued water maze acquisition. Behavioural Brain Research, 39(1), 29–51. 10.1016/0166-4328(90)90119-y

Selden, N., Robbins, T., & Everitt, B. (1990). Enhanced behavioral conditioning to context and impaired behavioral and neuroendocrine responses to conditioned stimuli following ceruleocortical noradrenergic lesions: Support for an attentional hypothesis of central noradrenergic function. The Journal of Neuroscience, 10(2), 531–539. 10.1523/jneurosci.10-02-00531.1990

Silveira, M. M., Murch, W. S., Clark, L., & Winstanley, C. A. (2016). Chronic atomoxetine treatment during adolescence does not influence decision-making on a rodent gambling task, but does modulate amphetamine’s effect on impulsive action in adulthood. Behavioural Pharmacology, 27(4), 350–363. 10.1097/FBP.0000000000000203

Smith, K. S., Bucci, D. J., Luikart, B. W., & Mahler, S. V. (2016). DREADDs: Use and Application in Behavioral Neuroscience. Behavioral Neuroscience, 130(2), 137–155. 10.1037/bne0000135

Su, Z., & Cohen, J. Y. (2022). Two types of locus coeruleus norepinephrine neurons drive reinforcement learning. Cold Spring Harbor Laboratory. 10.1101/2022.12.08.519670

Tervo, D. G. R., Proskurin, M., Manakov, M., Kabra, M., Vollmer, A., Branson, K., & Karpova, A. Y. (2014). Behavioral Variability through Stochastic Choice and Its Gating by Anterior Cingulate Cortex. Cell, 159(1), 21–32. 10.1016/j.cell.2014.08.037

Vazey, E. M., Moorman, D. E., & Aston-Jones, G. (2018). Phasic locus coeruleus activity regulates cortical encoding of salience information. Proceedings of the National Academy of Sciences of the United States of America, 115(40), E9439–E9448. 10.1073/pnas.1803716115

Vazquez, C. R., Becker, L. J., Kuo, C.-C., Cariello, S. A., Hamdan, A. N., Al-Hasani, R., Maloney, S. E., & McCall, J. G. (2025). Maternal separation disrupts noradrenergic control of adult coping behaviors. Neuropsychopharmacology, 1–12. 10.1038/s41386-025-02201-4

Velazquez-Sanchez, C., Muresan, L., Marti-Prats, L., & Belin, D. (2023). The development of compulsive coping behaviour is associated with a downregulation of Arc in a Locus Coeruleus neuronal ensemble. Neuropsychopharmacology, 48(4), 653–663. 10.1038/s41386-022-01522-y

Waterhouse, B. D., & Navarra, R. L. (2019). The locus coeruleus-norepinephrine system and sensory signal processing: A historical review and current perspectives. *Brain Research*, Behavioral Consequences of Noradrenergic Actions in Sensory Networks, 1709, 1–15. 10.1016/j.brainres.2018.08.032

Whelan, R., Conrod, P. J., Poline, J.-B., Lourdusamy, A., Banaschewski, T., Barker, G. J., Bellgrove, M. A., Büchel, C., Byrne, M., Cummins, T. D. R., Fauth-Bühler, M., Flor, H., Gallinat, J., Heinz, A., Ittermann, B., Mann, K., Martinot, J.-L., Lalor, E. C., Lathrop, M., … Garavan, H. (2012). Adolescent impulsivity phenotypes characterized by distinct brain networks. Nature Neuroscience, 15(6), 920–925. 10.1038/nn.3092

Williams, W. A., & Potenza, M. N. (2008). The neurobiology of impulse control disorders. Brazilian Journal of Psychiatry, 30, S24–S30. 10.1590/S1516-44462008005000003

Winstanley, C. A., & Hynes, T. J. (2021). Clueless about cues: The impact of reward-paired cues on decision making under uncertainty. *Current Opinion in Behavioral Sciences*, Value Based Decision-Making, 41, 167–174. 10.1016/j.cobeha.2021.07.001

