## Supplemental Figures for "Chemogenetic inhibition of the noradrenergic locus coeruleus promotes the development of risk-taking decisional strategies and selectively enhances motor impulsivity in females"

**Supplemental Figure 1: Behavioural effects of chemogenetic inhibition of LC wash out effectively**

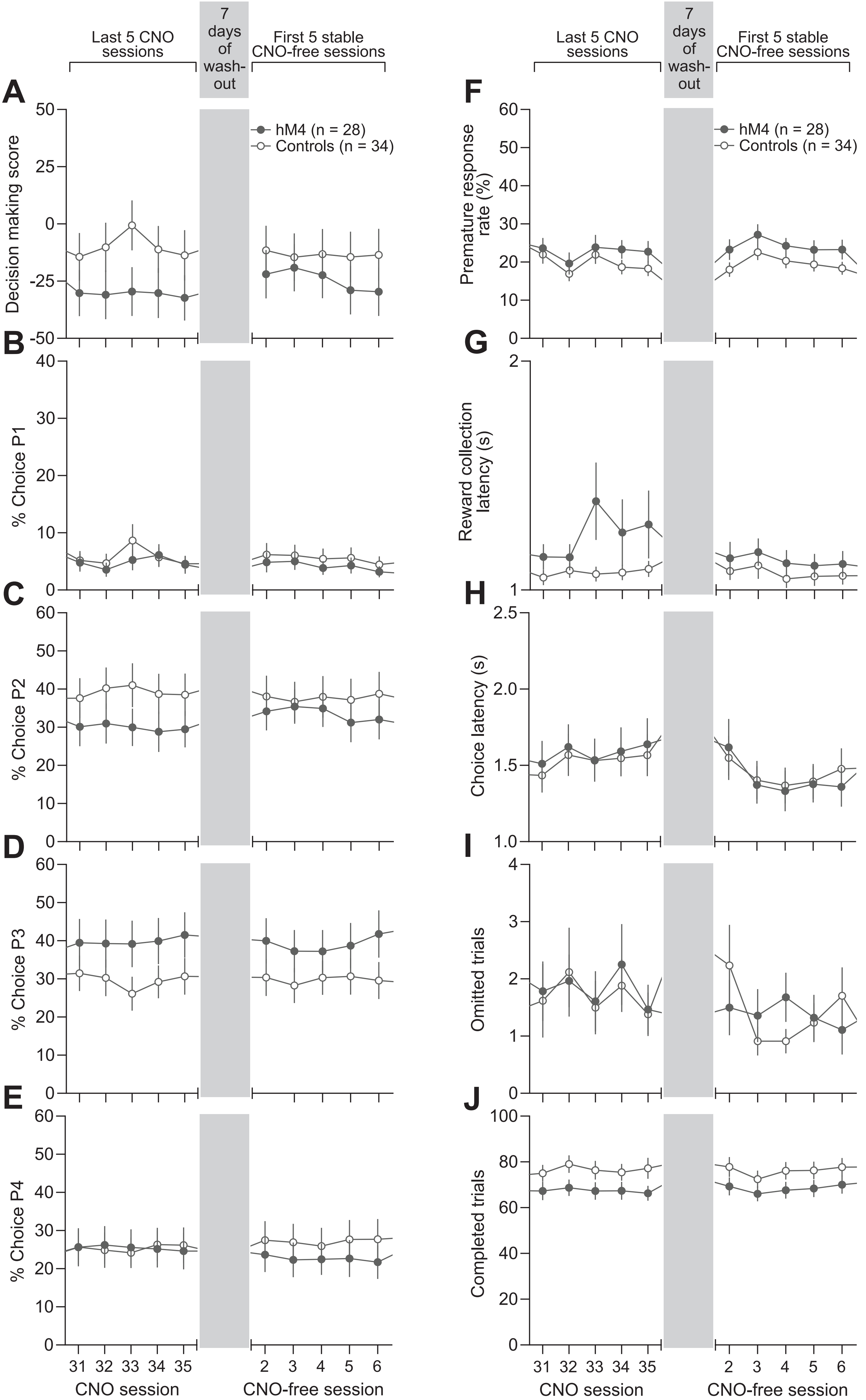

**Supplemental Figure 2: TG+ and TG- controls exhibit comparable behaviour on the crGT**

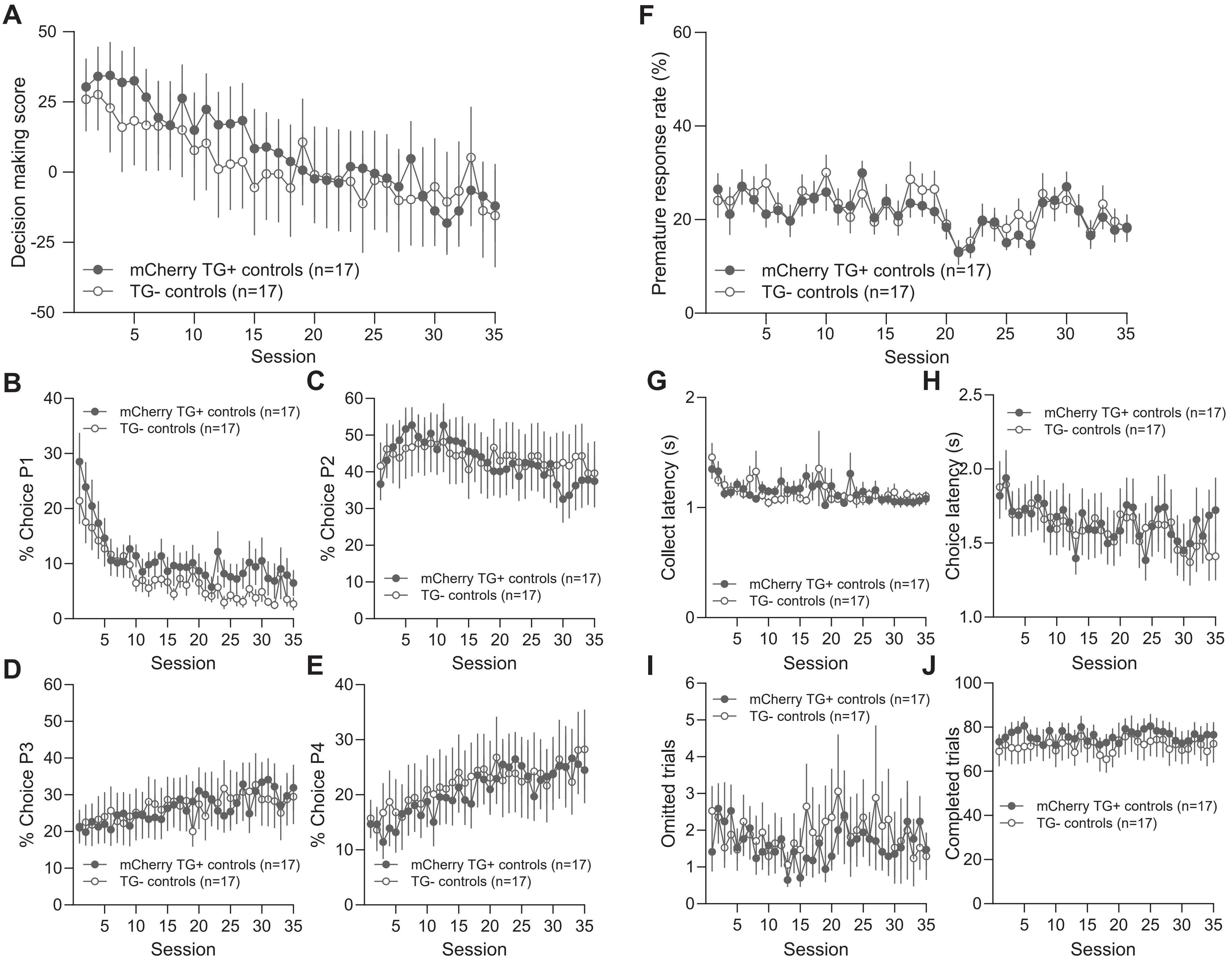
